# Massive gene amplification on a recently formed Drosophila Y chromosome

**DOI:** 10.1101/451005

**Authors:** Doris Bachtrog, Shivani Mahajan, Ryan Bracewell

## Abstract

Widespread loss of genes on the Y is considered a hallmark of sex chromosome differentiation. Here we show that the initial stages of Y evolution are driven by massive amplification of distinct classes of genes. The neo-Y chromosome of *Drosophila miranda* initially contained about 3000 protein-coding genes, but has gained over 3200 genes since its formation about 1.5 MY ago, primarily by tandem amplification of protein-coding genes ancestrally present on this chromosome. We show that distinct evolutionary processes may account for this drastic increase in gene number on the Y. Testis-specific and dosage sensitive genes appear to have amplified on the Y to increase male fitness. A distinct class of meiosis-related multi-copy Y genes independently co-amplified on the X, and their expansion is likely driven by conflicts over segregation. Co-amplified X/Y genes are highly expressed in testis, enriched for meiosis and RNAi functions, and are frequently targeted by small RNAs in testis. This suggests that their amplification is driven by X vs. Y antagonism for increased transmission, where sex chromosome drive suppression is likely mediated by sequence homology between the suppressor and distorter, through RNAi mechanism. Thus, our analysis suggests that newly emerged sex chromosomes are a battleground for sexual and meiotic conflict.

## Introduction

Sex chromosomes have originated multiple times from ordinary autosomes^1^. After suppression of recombination, the proto-X and proto-Y chromosomes follow separate evolutionary trajectories and differentiate^2^. The complete lack of recombination on Y chromosomes renders natural selection inefficient, and Y evolution is characterized by a loss of the majority of its ancestral genes^3^ while the X is thought to maintain most of them. Indeed, old Y chromosomes of various species contain only a few functional genes^4,5^, and Y chromosomes of many taxa (but not all) have instead accumulated massive amounts of repetitive DNA, including transposable elements (TEs) and satellite DNA^6^. In some lineages, the Y chromosome is lost entirely^7^.

Studies of Y chromosomes are often hindered by a lack of high-quality reference sequences, due to the technical challenges of assembling repetitive regions. To date, the Y chromosomes of only a handful of mammalian species have been fully sequenced^8,9^ and no high-quality sequences of young Y chromosomes that have already accumulated a substantial amount of repetitive DNA have been examined.

*Drosophila miranda* has a pair of recently formed neo-sex chromosomes that originated ~1.5MY ago after its split from its closely related sister species *D. pseudoobscura*, and has served as a model to study the initiation of sex chromosome differentiation^10^. The neo-sex chromosomes of *D. miranda* were formed by the fusion of a former autosome (chr3 of the *pseudoobscura* group^11^) with the ancestral, degenerate Y chromosome of this clade^12^. The neo-X and neo-Y chromosome are still homologous over much of their length, with 98.5% sequence identity at homologous regions^10^. A previous genomic analyses using Illumina short reads confirmed the notion that genes on the Y are rapidly lost^13^. About ^1^/_3_ of the roughly 3000 genes ancestrally present on the neo-Y were found to be pseudogenized, and over 150 genes were entirely missing^13^. However, the high level of sequence similarity between the neo-X and neo-Y chromosome, yet drastic accumulation of repeats on the neo-Y, prevented assembling the Y chromosome using short-read data.

We recently generated a high-quality sequence assembly of the neo-Y chromosome of *D. miranda* using single-molecule sequencing and chromatin conformation capture, and used extensive BAC clone sequencing and optical mapping data to confirm that our assembly is of high accuracy^14^. Intriguingly, instead of simply shrinking, our assembly revealed that the young neo-Y chromosome dramatically increased in size relative to the neo-X by roughly 3-fold. We assembled 110.5 Mb of the fused ancestral Y and neo-Y chromosome (Y/neo-Y sequence), and 25.3 Mb of the neo-X^14^. Most of this size increase is driven by massive accumulation of repetitive sequences —in particular transposable elements— which comprise over 50% of the neo-Y derived sequence^14^. Here, we carefully annotate the neo-sex chromosomes using transcriptomes from multiple tissues and small RNA profiles, to study the evolution of gene content on this recently formed neo-Y chromosome.

## Results

### A catalogue of genes on the neo-Y

With a comprehensive high-quality reference sequence of the neo-Y chromosome of *D. miranda*, we systematically catalogued its genes. Comparison of the neo-sex chromosome gene content with that of *D. pseudoobscura*, a close relative where this chromosome pair is autosomal, allows us to infer the ancestral gene complement and reconstruct the evolutionary history of gene gains and losses along the neo-sex chromosomes (Figure 1A). Note that the ancestral Y chromosome of *D. pseudoobscura* is not assembled and contains no annotated protein-coding genes, and our analysis focuses on neo-sex linked genes (i.e. genes present on chr3 of *D. pseudoobscura*). Annotation of neo-Y linked genes is a challenging task, for several reasons. Genes on the neo-Y are embedded in highly repetitive sequences, and introns often dramatically increase in size due to TE insertions^15^. Neo-Y genes (or pseudogenes) are also often truncated or have premature stop-codons^13,16^. Automated annotation thus often resulted in fragmented, split or missing gene models on the neo-Y (see Methods for details), and we used extensive manual curation to validate and correct our gene models (see Methods). In particular, we bioinformatically identified and manually examined (and if necessary corrected) all neo-Y genes that were not simple 1:1 orthologs between species and neo-sex chromosomes. In total, we identified 6,448 genes on the neo-Y, and 3,253 genes on the neo-X, compared to 3,087 genes on the ancestral autosome that gave rise to the neo-sex chromosome. Thus, contrary to the paradigm that Y chromosomes undergo chromosome-wide degeneration, our analysis reveals a dramatic increase in the number of annotated genes on the neo-Y, compared to its ancestral gene complement, or that of its former homolog, the neo-X.

**Figure 1.**
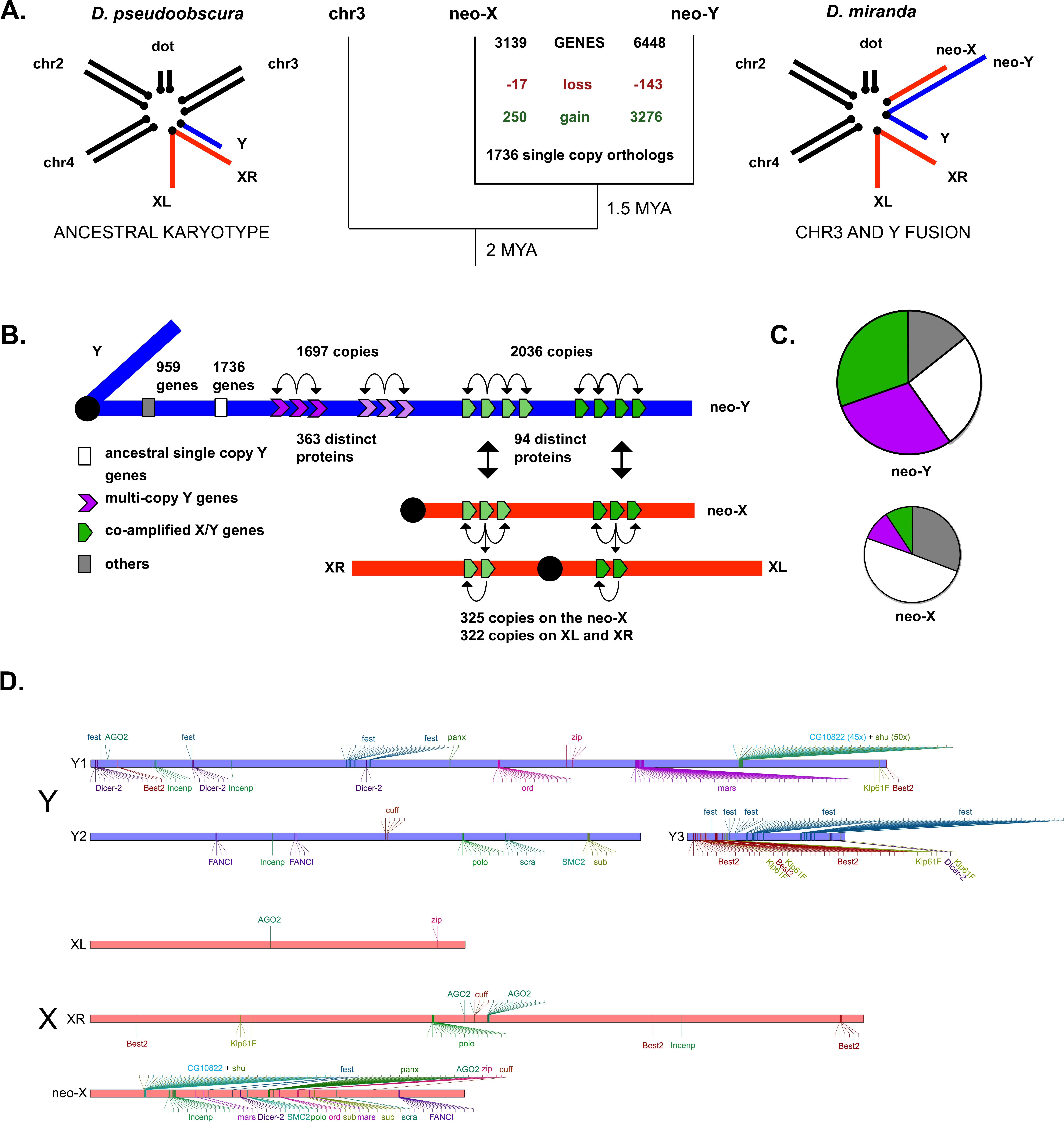
Gene content evolution of newly formed sex chromosomes. **A.** Karyotype and gene content evolution on the neo-sex chromosomes of *D. miranda*. Shown are the karyotype of *D. miranda*, and its close relative *D. pseudoobscura*, from which it diverged about 2MY ago. In *D. miranda*, the fusion of an autosome (chromosome 3) with the Y chromosome created the neo-sex chromosomes, about 1.5 MY ago. Shown along the tree are numbers of gene amplifications (in green) and gene losses (in red) of genes ancestrally present on chr3, on the neo-X and neo-Y chromosomes, assuming parsimony. X chromosomes are shown in red, Y chromosomes in blue, and autosomes in black. **B**. Schematic representation of gene content of the *D. miranda* neo-Y/Y chromosome. The *D. miranda* neo-Y/Y contains different types of genes: our annotation contains 1736 ancestral single-copy genes; 1697 multi-copy Y genes that are derived from 363 distinct proteins that were ancestrally present on the neo-Y, and 2036 genes that are derived from 94 distinct proteins that were ancestrally present on the autosome that formed the neo-sex chromosome and that co-amplified on both the X/neo-X and Y/neo-Y. “Others” refers to genes not present or mapping to an unknown location in *D. pseudoobscura* (446 genes), or genes with complex mapping (513 genes; see Methods). **C.** Pie charts show the assignment of genes on the neo-Y/Y or neo-X to these different categories (using the same color scheme as in panel B), with the size of the pie scaled by the number of genes on the neo-Y/Y or neo-X. **D.** Co-amplified X/Y genes typically exists as tandem repeats on the X and the Y chromosomes. Shown are a subset of 18 co-amplified X/Y gene families with meiosis and siRNA functions.

Overall, we detect 1,736 ancestral single-copy orthologs between the neo-sex chromosomes, i.e. roughly 56% of genes ancestrally present show a simple 1:1 relationship between species and the neo-X and neo-Y chromosome. Furthermore, genes are degenerating on the non-recombining neo-Y. We find 143 genes that are located on chr3 in *D. pseudoobscura* and the neo-X of *D. miranda*, but are missing from our neo-Y annotation, and we fail to detect a homolog on the neo-Y by BLAST. Thus, about 5% of genes that were ancestrally present are now completely absent on the neo-Y. On the other hand, only 17 genes (0.5% of genes ancestrally present) are absent from the neo-X but found on the neo-Y, a rate of gene loss comparable to autosomes and the ancestral X (**Table S1**). Thus, the neo-Y is indeed losing its ancestral genes at a high rate, consistent with theoretical expectation^3,17^ and empirical observations of gene poor ancestral Y chromosomes^4,5,8,9^.

Intriguingly, however, for 457 unique single-copy *D. pseudoobscura* protein-coding genes, we find multiple copies in our neo-Y chromosome annotation of *D. miranda* (which were all verified by manual inspection of BLAST and nucmer alignments^18^, see Methods for details; and also confirmed by Illumina read depth analysis, see **Figure S1**). Genes with multiple copies on the Y/neo-Y fall into two groups. 363 unique protein-coding genes of *D. pseudoobscura* are also single-copy (or missing) on the X/neo-X of *D. miranda*, but are amplified on the neo-Y (resulting in a total of 1,697 Y-linked gene copies; two of these genes were gained from chr2; **Table S2, Figure S2, S3**). The remaining 94 unique protein-coding genes of *D. pseudoobscura* that have amplified on the Y/neo-Y, surprisingly, have also co-amplified on the original X and neo-X chromosome of *D. miranda* (and harbor a total of 2,036 Y/neo-Y-linked gene copies and 650 copies on the X/neo-X; **Table S3**, Figure 1D). Most of the genes that co-amplified on the X and Y chromosome of *D. miranda* were ancestrally present on the autosome that formed the neo-sex chromosomes (i.e. chr3 in *D. pseudoobscura*), but some were also gained from other chromosomes (4 genes from chr2, 1 gene from chr4, and 14 genes from the ancestral X; **Figure S3**). Thus, genes on the Y/neo-Y of *D. miranda* fall into distinct categories (Figure 1B, **Figure S3**), and we refer to them as single-copy Y genes, multi-copy Y genes (which are single-copy on the X/neo-X), and co-amplified X/Y genes (genes that have amplified on both the X and the Y chromosome). Genes whose ancestral location could not be determined, or genes with more complex evolutionary histories were not further analyzed (see Methods).

### Properties of amplified Y genes

Many amplified gene copies – on both the X/neo-X and the Y/neo-Y chromosome - are fragmented, and some have premature stop codons or frame shift mutations (**Table S2, S3**). We find full-length copies for 786 amplified Y genes (46%), 776 co-amplified Y-genes (38%), and 300 co-amplified X genes (46%). Thus, even if ignoring partial gene copies (which may nevertheless have function as non-coding transcripts), we still find considerably more genes on the neo-Y compared to the neo-X or the ancestral autosome that formed the neo-sex chromosomes. Genes with truncated coding regions are less likely to produce functional proteins and may thus be pseudogenes. However, many of these amplified gene copies may instead encode functional RNAs (for example, they may be involved in RNA induced silencing, as suggested by our analysis below), and thus both full-length and fragmented copies could influence organismal fitness, if expressed. Indeed, transcriptome analysis (using only uniquely mapping RNA-seq reads) shows that most individual gene copies of amplified gene families on the Y/neo-Y are expressed in male tissues, both for partial genes and full-length transcripts. We detect expression of 71% of individual copies among multi-copy Y genes, and 94% for co-amplified X/Y genes (**Table S4**). This is consistent with many gene copies on the neo-Y indeed being functional, either as a protein or as a functional RNA.

How do genes amplify on the sex chromosomes? The majority of multi-copy Y genes, or co-amplified X and Y genes are found in gene clusters (Figure 1D, **Figure S2; Table S5, S6**). In particular, we find that 89% of multi-copy Y genes are located near (within 100 kb) other copies of the same gene family, and 80% of co-amplified X and 87% of co-amplified Y genes (Figure 1D, **Figure S2; Table S5, S6**). Clustering of gene families in tandem arrays suggests that non-allelic recombination is a main factor driving gene amplification on sex chromosomes. Additionally, phylogenetic analysis reveals that individual copies of co-amplified gene families typically cluster by chromosome, indicating independent amplification on the X and Y chromosome, and confirming a lack of recombination between the neo-X and neo-Y (**Figure S4**). Multi-copy genes often show dynamic copy number evolution between individuals^19^. To test for variation in copy number of amplified Y/neo-Y genes in natural populations of *D. miranda*, we generated Y-chromosome replacement lines by backcrossing Y chromosomes from different locations into the same genetic background (**Table S7, Figure S5**; see Methods). This strategy avoids confounding variation at X and autosomal regions, and Y copy number polymorphisms were estimated based on Illumina read coverage (see Methods). Overall, we find relatively little variation in copy number for both multi-copy Y and co-amplified Y genes among different neo-Y chromosomes (**Figure S6**). Low copy number variation is consistent with reduced levels of single-nucleotide diversity on the *D. miranda* neo-Y chromosome (π=0.01%; i.e. 30-fold lower than typical levels of variation in this species^20^), due to a recent selective sweep that completely eliminated all standing variation a few thousand years ago^20^.

## Discussion

### Different evolutionary processes may cause amplification of neo-Y genes

What may drive massive gene amplification on the neo-Y chromosome? Y chromosomes are subject to unique evolutionary forces: they lack recombination, show male-limited inheritance, and compete with the X over transmission to the next generation^3,21^. Indeed, our functional genomic analysis suggests that different processes appear to trigger gene family expansion of multi-copy Y genes versus co-amplified X/Y genes (Figure 2). Repetitive sequences, and in particular transposable elements are accumulating on the Y, and its high repeat content makes the Y chromosome particularly prone to accumulate multi-copy genes for multiple reasons (Figure 2A). On one hand, repetitive sequences can provide a substrate for non-allelic homologous recombination and thereby promote gene family expansion^22^. Indeed, we find several cases where repeats flank gene duplications on both the X and Y chromosome, and may have contributed to their origination (**Figure S7**). Additionally, spreading of heterochromatin from repeats globally dampens expression of neo-Y genes^23^, and multi-copy gene families may simply be more tolerated on the neo-Y (although many individual gene copies are transcribed, **Table S4**).

**Figure 2.**
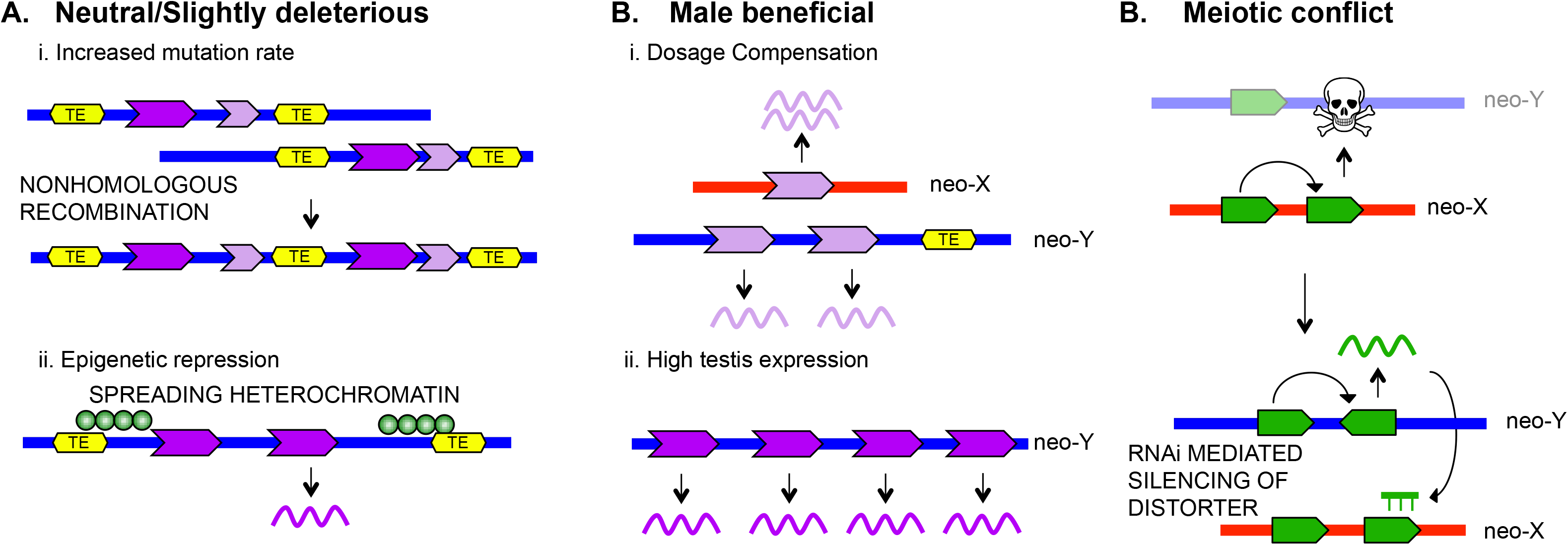
Distinct evolutionary processes may drive the accumulation of multi-copy Y genes, or co-amplified X and Y genes. **A.** Amplified Y genes may have no fitness benefits or be slightly deleterious. Repeats on the neo-Y can provide a substrate for non-allelic homologous recombination and promote gene family expansion (i). Gene duplicates may be silenced by spreading heterochromatin on the neo-Y, and thus less deleterious (ii). **B.** Multi-copy Y genes may provide fitness benefits to males, either through compensating for reduced gene dose of neo-Y genes (i) or by contributing to male fertility (ii). C. Co-amplified X/Y genes may be involved in an intergenomic conflict over segregation, and invoke the RNAi pathway to trigger silencing of meiotic drivers.

Gene family expansions on the Y can also be beneficial for males (Figure 2B). Global transcription is lower from the neo-Y chromosome of *D. miranda*^24^, and drives the evolution of dosage compensation of homologous neo-X genes^25^. In Drosophila, dosage compensation is achieved by transcriptional up-regulation of X-linked genes in males^26^. *D. miranda* has evolved only partial dosage compensation of its neo-X chromosome^25,27^, and gene amplification may help compensate for reduced gene dose of neo-Y genes, especially if their neo-X homologs are not yet dosage compensated. Additionally, Y chromosomes are transmitted from father to son, and are thus an ideal location for genes that specifically enhance male fitness^28^. Y chromosomes of several species, including humans, have been shown to contain multi-copy gene families that are expressed in testis and contribute to male fertility^29–31^.

Gene amplification on the Y could also be a signature of intragenomic conflicts (Figure 2C). Y chromosomes compete with the X over transmission to the next generation^32,33^, and sex chromosomes may try to cheat fair meiosis to bias their representation in functional sperm (i.e. meiotic drive). Meiotic drive on sex chromosomes, however, reduces fertility and distorts population sex ratios^32^, and creates strong selective pressure to evolve suppressors to silence selfish drivers. Suppression of sex chromosome drive could be mediated by sequence homology between the suppressor and distorter, through RNAi mechanisms, and could result in co-amplification of genes on the X and Y chromosome. The RNAi pathway has been implicated to mediate suppression of sex chromosome drive in Drosophila^34–36^.

### Amplification of multi-copy neo-Y genes may increase male fitness

We find a handful of multi-copy Y gene families that have dozens of gene copies on the Y (six gene families have more than 30 copies, and 14 gene families have >15 copies; Figure 3A, B, **Figure S8**), while the vast majority of multi-copy Y gene families only have a few copies (90% of multi-copy Y gene families have fewer than 4 copies). Dosage compensation counters ploidy differences of X-linked genes in males vs. females (i.e. one vs. two copies), and thus may contribute to amplification of multi-copy Y genes with few copies (i.e. 2-4 copies on the Y), while testis-expressed multi-copy Y genes often contain dozens of gene copies^29–31,37^. Gene expression and chromatin analysis support that different evolutionary forces may contribute to the accumulation of low versus high-copy number multi-copy Y gene families.

**Figure 3.**
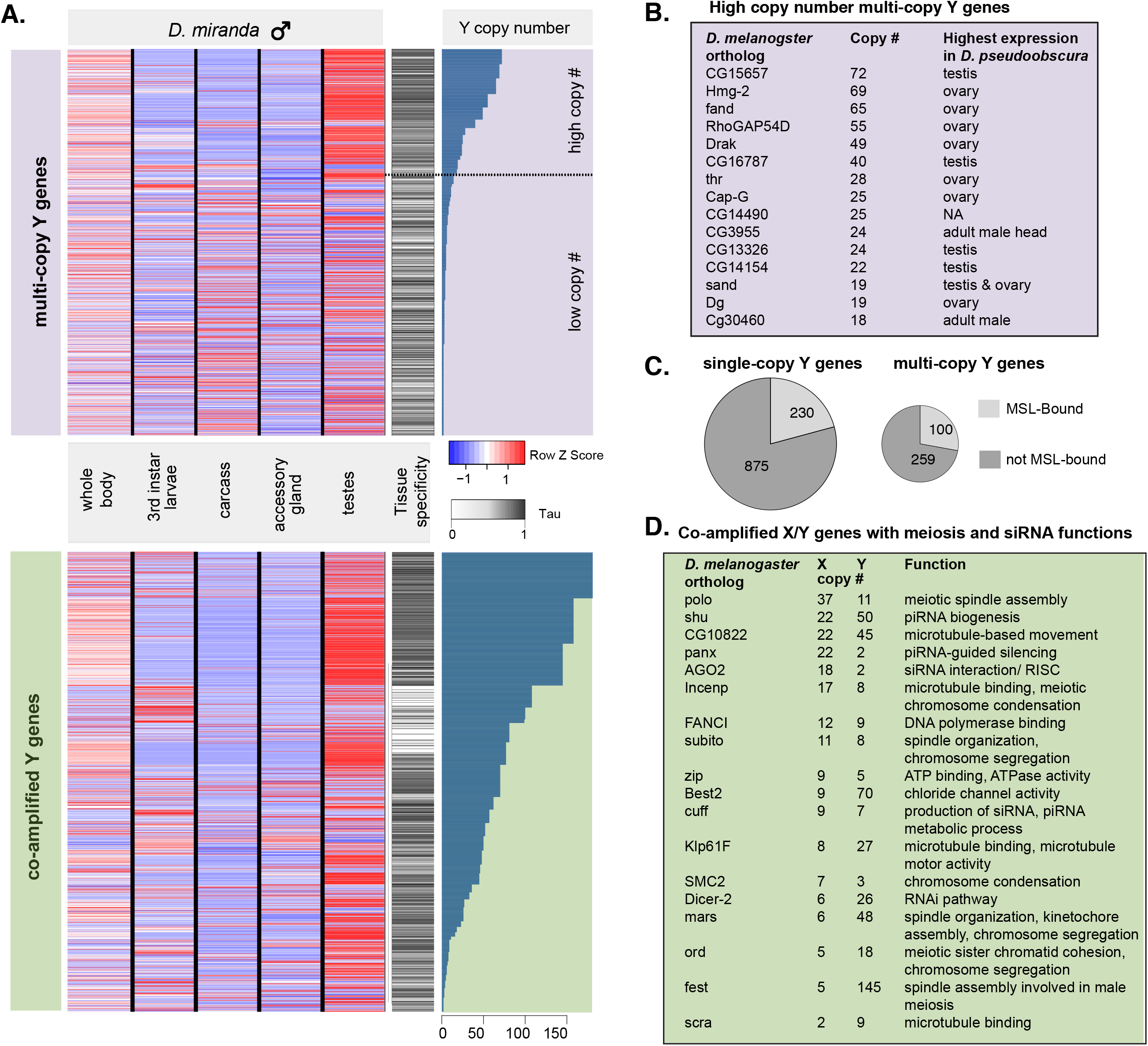
Characterization of ampliconic Y genes. **A**. Expression of multi-copy Y genes (top, purple shading) and co-amplified Y genes (bottom, green shading) in different male *D. miranda* samples, and tissue-specificity index. Genes are sorted by their copy number on the Y, and the tissue-specificity index, is calculated as described in Methods. Multi-copy Y genes with high copy number are primarily expressed in testis, while gene families with low copy number are expressed in multiple tissues. The Z-Scores are calculated by scaling the rows to have 0 median and a standard deviation of 1. The data shown in panel A are presented in **Data Supplement 6**. **B.** List of top amplified multi-copy Y genes (i.e. high copy number Y genes), their copy number on the Y, and tissue of highest expression on *D. pseudoobscura* (as a proxy for ancestral expression). **C.** Dosage compensation status of neo-X homologs of single-copy (left) and multi-copy (right) Y genes. Shown are the relative numbers of neo-X genes that are bound by the MSL-complex (i.e. that are dosage compensated), and those not bound (i.e. not dosage compensated). MSL-binding data were generated for male *D. miranda* larvae (28). Genes with multiple copies on the Y are less likely to be dosage compensated on the X. The data in panel C are presented in **Data Supplement 7**. **D.** List of top co-amplified X/ Y genes with well-characterized roles in meiosis or siRNA, their copy number, and inferred function in D. melanogaster.

Multi-copy Y gene families with a high copy number (i.e. >15 copies) are expressed almost exclusively in testis (Figure 3A, **Figure S8, S9**), mimicking patterns of gene family amplification of male fertility genes found in other species^4,29,31,38,39^. Their neo-X homologs, in contrast, are expressed predominantly in ovaries (**Figure S8, S9**). Gene expression profiles in *D. pseudoobscura* suggests that these genes were ancestrally highly expressed in testis and/or ovaries (Figure 3B), and sex-linkage may have enabled neo-Y and neo-X gametologs to specialize in their putative male- and female specific function, respectively^28^. Most multi-copy Y gene families, in contrast, only have few copies and are ubiquitously expressed (Figure 3A). Consistent with gene dosage contributing to increased copy number on the Y, the neo-X homologs of multi-copy Y genes are less likely to be dosage compensated compared to single-copy Y genes (Figure 3C). In particular, male Drosophila achieve dosage compensation by recruiting the MSL-complex to their hemizygous X chromosome^26^, and neo-X homologs of multi-copy neo-Y genes are less likely to be targeted by the MSL complex in male *D. miranda* larvae^27^ than the neo-X homologs of single-copy neo-Y genes (Figure 3C, p-value Fisher’s exact test = 0.007). This suggests that many multi-copy Y genes are dosage sensitive, and additional gene copies on the Y may contribute to dosage compensation. Truncated neo-Y genes are less likely to produce functional proteins and thus alleviate gene dose deficiencies. Despite having many fewer copies on the Y on average (average 3 copies per gene), the low copy number multi-copy Y genes have a similar fraction of genes with at least two full-length copies (roughly half) as high-copy number multi-copy Y genes (average 26 copies per gene), or co-amplified X/Y genes (22 copies per gene; see **Table S2, S3**). Genes that co-amplify on the X and Y chromosome, on the other hand, show testis-biased expression, independent of copy number (Figure 3A). Gene ontology (GO) analysis found no significant enrichment of gene annotations among multi-copy Y genes, consistent with a broad category of (possibly dosage sensitive) genes amplifying on the Y. Overall, our analysis is consistent with a considerable fraction of multi-copy Y genes having an important function in males, as supported by tissue-specific expression, or patterns of MSL-binding.

### X/Y co-amplified genes suggest ongoing conflict over sex chromosome transmission

Functional enrichment analysis (Figure 4A, B), gene expression patterns and small RNA profiles (Figure 5) suggest that fundamentally different forces drive co-amplification of genes on the X and Y chromosome. Overall, we identify 2683 co-amplified genes on the neo-sex chromosomes of *D. miranda* (2036 genes amplified on the Y/neo-Y, and 650 on the X/neo-X). Co-amplified X/Y genes belong to 94 distinct proteins that were ancestrally single-copy on the autosome that formed the neo-sex chromosomes of *D. miranda* (that is, these genes are single-copy on chr3 of *D. pseudoobscura*), and phylogenetic analysis confirms their independent amplification on the X and Y (**Figure S4**). Co-amplified X and Y-linked gene copies are typically both highly expressed in testis (Figure 3A, Figure 5B; **Figure S8, S9**). Testis expression of co-amplified X-linked genes is unusual, as testis-genes in *Drosophila* normally avoid the X chromosome^40–43^, but can be understood under intragenomic conflict models^21,34,35,44,45^. In particular, an X-linked gene involved in chromosome segregation may evolve a duplicate that acquires the ability to incapacitate Y-bearing sperm (Figure 2C). Invasion of this segregation distorter skews the population sex ratio and creates a selective advantage to evolve a Y-linked suppressor that is resistant to the distorter. Suppression may be achieved at the molecular level by increased copy number of the wildtype function or by inactivation of X-linked drivers using RNAi^34–36^. If both driver and suppressor are dosage-sensitive, they would undergo iterated cycles of expansion, resulting in rapid co-amplification of both driver and suppressor on the X and Y chromosome^32^.

**Figure 4.**
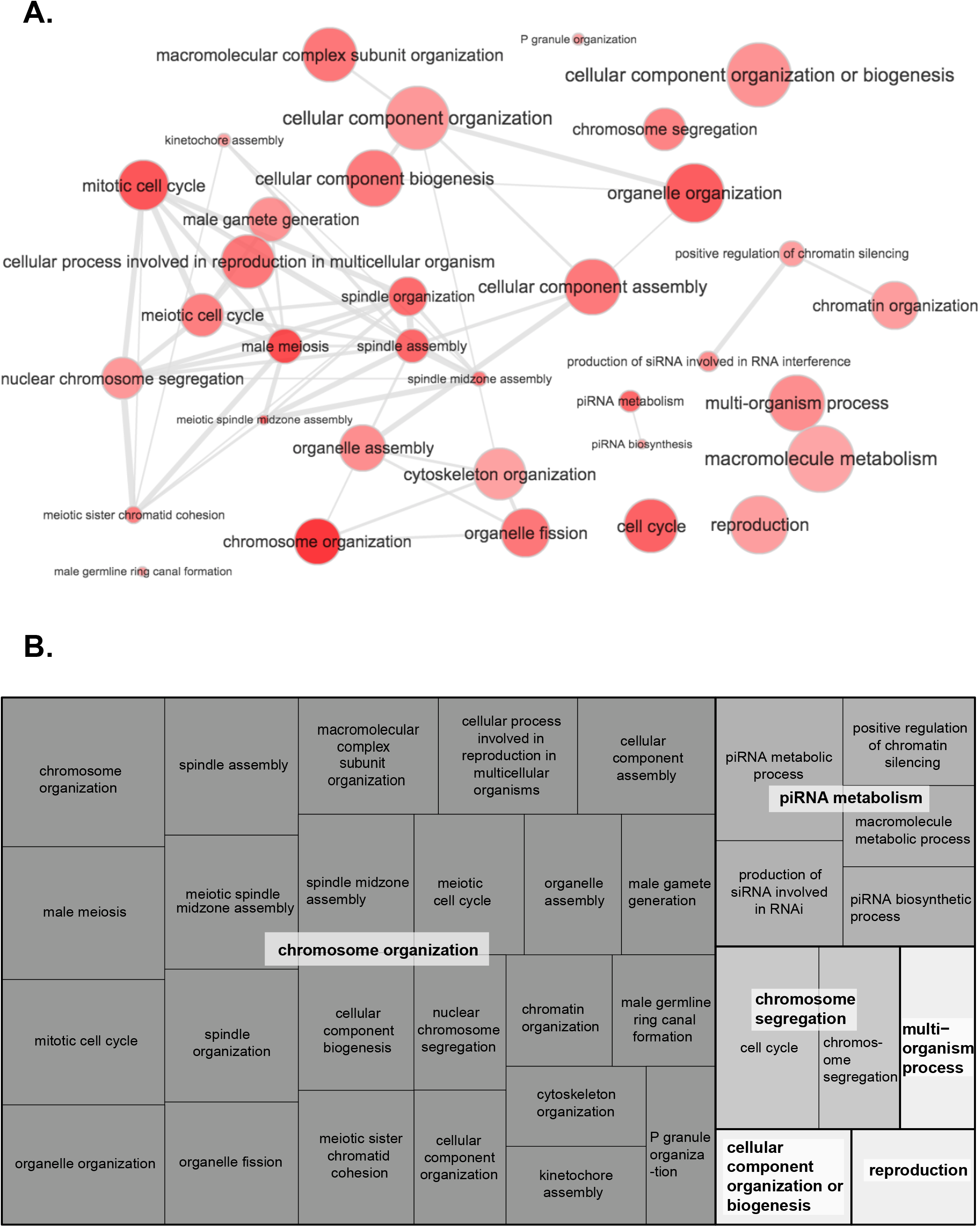
Co-amplified X/Y genes are enriched for meiosis-related and RNAi functions. **A.** “Interactive graph” view of enriched GO terms. Bubble color indicates the *p*-value; bubble size indicates the frequency of the GO term in the underlying GO database. Highly similar GO terms are linked by edges in the graph, where the line width indicates the degree of similarity. **B.** “TreeMap” view of enriched GO terms. Each rectangle is a single cluster representative. The representatives are joined into ‘superclusters’ of loosely related terms, visualized with different shades of gray. Size of the rectangles reflect the *p*-value of the GO term.

**Figure 5.**
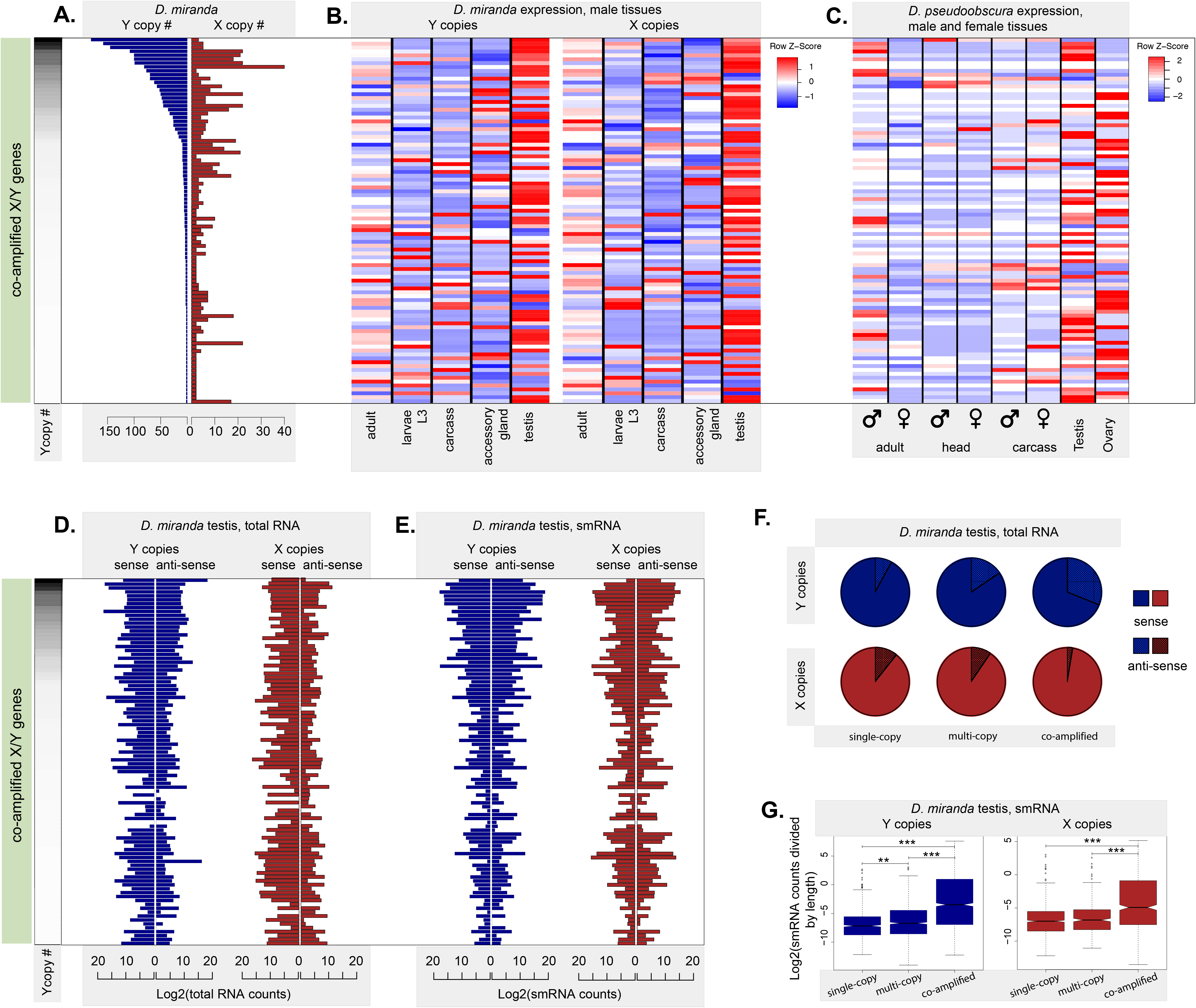
Co-amplified X/Y gene families produce anti-sense transcripts and small RNAs in testis. The 94 co-amplified gene families are sorted by copy number on the Y in panels A-E. **A.** Copy numbers on the Y and X chromosomes for co-amplified gene families in *D. miranda*. The data shown in panel A are presented in **Data Supplement 8**. **B.** Tissue expression patterns for co-amplified X and Y genes in *D. miranda* male tissues. Co-amplified X/Y genes are highly expressed in testis. The Z-Scores are calculated by scaling the rows to have 0 mean and standard deviation 1. The data shown in panel B are presented in **Data Supplement 9**. **C.** Tissue expression patterns of homologos of co-amplified X/Y genes in *D. pseudoobscura.* Homologs of co-amplified X/Y genes are highly expressed in testis and ovaries, suggesting an ancestral function in gametogenesis. The Z-Scores are calculated by scaling rows to have 0 mean and standard deviation 1. The data shown in panel C are presented in **Data Supplement 10**. **D.** Sense- and anti-sense transcription of total RNA for co-amplified X and Y genes in *D. miranda* testis. Shown are testis total RNA counts derived from sense and antisense transcripts. The data shown in panel D are presented in **Data Supplement 11**. **E.** Sense- and anti-sense counts of small RNA for co-amplified X and Y genes in *D. miranda* testis. Shown are testis small RNA counts derived from sense and antisense transcripts. The data shown in panel E are presented in **Data Supplement 12**. **F.** Fraction of sense and anti-sense transcripts produced for different categories of genes on the X/neo-X and Y/neo-Y chromosome (i.e. ancestral single-copy Y and X genes; multi-copy Y/neo-Y genes and their neo-X gametologs; genes co-amplified on the Y/neo-Y and X/neo-X). The data shown in panel E are presented in **Data Supplement 13**. **G.** Enrichment of small RNAs mapping to co-amplified X and Y genes. Shown are testis small RNA counts (normalized by total gene length for all copies of a gene family) for different categories of genes on the X/neo-X and Y/neo-Y chromosome (i.e. ancestral single-copy Y and X genes; multi-copy Y/neo-Y genes and their neo-X gametologs; genes co-amplified on the Y/neo-Y and X/neo-X). The Wilcoxon test p-value significance is denoted by asterisks. The upper whisker and lower whisker in the boxplots show the 75th percentile + 1.5 times the interquartile range and the 25th percentile - 1.5 times the inter-quartile range, respectively. The data shown in panel G are presented in **Data Supplement 14**.

Consistent with a model of ongoing conflicts over chromosome segregation driving co-amplification of X/Y genes, we find that many of the most highly co-amplified genes have well-characterized functions in meiosis (Figure 3D, **Table S3**), and are ancestrally expressed in gonads (using gene expression data from *D. pseudoobscura* as a proxy for ancestral expression; Figure 5C, **Figure S10**). Gene ontology (GO) analysis reveals that co-amplified X/Y genes are significantly overrepresented in biological processes associated with meiosis and chromosome segregation (Figure 4A, 3B). In particular, multi-copy Y genes are significantly enriched for GO categories including “nuclear division”, “spindle assembly”, “meiotic spindle midzone assembly”, “DNA packaging”, “chromosome segregation”, or “male gamete generation” (see **Table S3**). Among the most highly co-amplified X/Y genes are well-studied genes with important function in meiosis, including *wurstfest* (145 copies on the Y and 5 on the X), a gene involved in spindle assembly in male meiosis I; *mars* (48 Y-linked copies and 6 X-linked copies), a gene involved in kinetochore assembly and chromosome segregation, *orientation disruptor* (18 Y-linked copies and 5 X-linked copies), a chromosome-localized protein required for meiotic sister chromatid cohesion, or *Subito* (8 Y-linked copies and 11 X-linked copies), a gene required for spindle organization and chromosome segregation in meiosis (Figure 3D, Figure 4, see **Table S3** for additional genes). These important meiosis genes are typically single-copy and highly conserved across insects, but highly co-amplified on the recently evolved *D. miranda* X and Y chromosome.

### Possible involvement of RNAi in sex chromosome drive

Additionally, our GO analysis reveals a significant overrepresentation of co-amplified X/Y genes associated with piRNA metabolism and the generation of small RNA’s (Figure 4A, 3B). Again, this is expected under recurring sex chromosome drive where silencing of distorters is achieved by RNAi, since compromising the small RNA pathway would release previously silenced drive systems^36^. Noteworthy genes in the RNAi pathway that are typically single-copy in insects but co-amplified on the X and Y of *D. miranda* include *Dicer-2* (26 Y- and 6 X-linked copies), a double-stranded RNA-specific endonuclease that cuts long double-stranded RNA into siRNAs, *cutoff* (7 Y- and 9 X-linked copies), a gene involved in transcription of piRNA clusters, or *shutdown* (50 Y- and 22 X-linked copies), a co-chaperone necessary for piRNA biogenesis (Figure 3D, Figure 4, **Table S3**). Thus, functional enrichment supports a model of meiotic conflict driving co-amplification of X/Y genes.

We gathered stranded RNA-seq and small RNA profiles from wildtype *D. miranda* testis, to obtain insights into the molecular mechanism of putative sex chromosome drive. Consistent with a model of meiotic drive and suppression through RNAi mechanisms causing co-amplification of X/Y genes, we detect both sense and antisense transcripts and small RNA’s derived from the vast majority of co-amplified X/Y genes (Figure 5D-G, Figure 6). Globally, we find that co-amplified Y genes show significantly higher levels of anti-sense transcription and small RNA production than single-copy Y genes, or multi-copy Y genes (Figure 5D,F; Wilcoxon test p-value < 10^−16^, **Table S8**). Likewise, small RNA levels are higher for co-amplified X linked genes, compared to single-copy X genes, or X homologs of multi-copy Y genes (Figure 5E,G; Wilcoxon test p-value < 10^−16^, **Table S8**). Anti-sense transcription of many co-amplified X/Y genes suggests that they may function not as proteins, but instead as functional RNA by generating double-stranded RNA and triggering the RNAi silencing pathway. Targeting of co-amplified X/Y genes by small RNA’s in testis demonstrates that small RNA production is not simply a consequence of the repeat-rich environment of the neo-Y but instead a property of co-amplified X/Y genes. Overall, our data are consistent with sex chromosome drive having repeatedly led to characteristic patterns of gene amplification of homologous genes on both the X and the Y chromosomes that are targeted by small RNAs (Figure 6).

**Figure 6.**
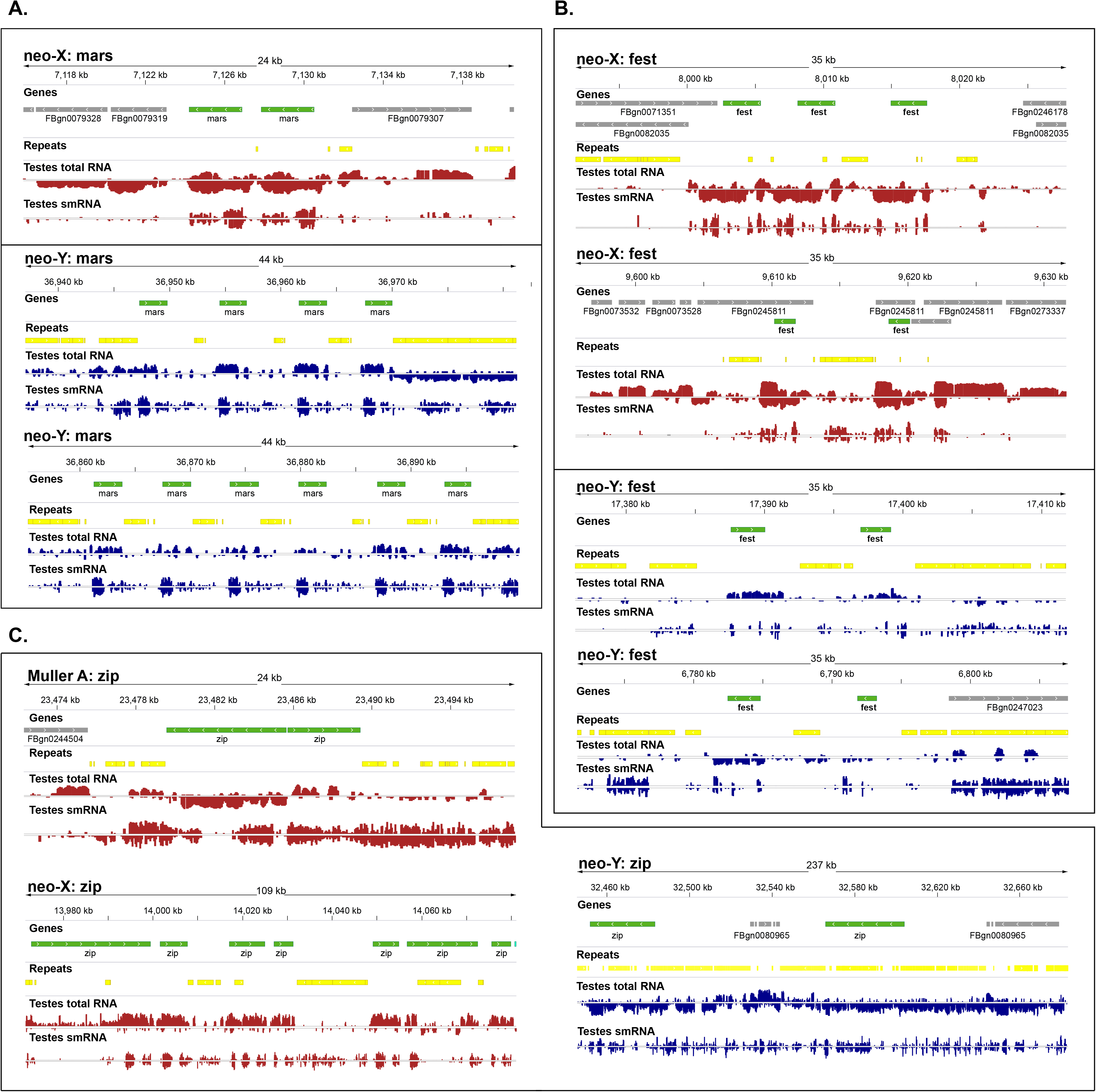
Examples of co-amplified X and Y genes. Shown are the genomic architecture of co-amplified gene families on the neo-X and neo-Y (repetitive regions are displayed in yellow, and co-amplified genes in green, other genes in gray), and expression profiles from testis (stranded RNA-seq and small RNA profiles in red for the neo-X and blue for the neo-Y). For each gene (**A.** *fest*; **B.** *mars*; **C.** *zip*), only a representative subset of copies is shown.

## Conclusions

Contrary to the paradigm that Y chromosomes undergo global degeneration, we document a high rate of gene gain on the recently formed neo-Y chromosome of *D. miranda*, mainly through amplification of genes that were ancestrally present on the autosome (chr3) that became the neo-Y. Our comparative genomic analysis reveals different types of amplified Y genes, and we show that their acquisition likely is driven by different selective pressures. Multi-copy genes exclusive to the Y presumably increase male fitness, while genes that are co-amplified on the X and Y likely reflect intragenomic conflict. Multi-copy Y genes come in two flavors, and our analysis suggests that they are either selected and amplifying on the Y because of their testis-specific function, or to compensate for gene dosage deficiencies. Genes with testis-biased expression often have dozens of copies on the Y, and their neo-X homologs are often expressed in ovaries, and sex linkage may have allowed these former homologs to specialize in their sex-specific roles^36^. Ubiquitously expressed housekeeping genes also duplicate on the Y, possibly to mitigate gene dose deficiencies of partially silenced neo-Y genes; these genes are present at a much lower copy number, and are targeted less often by the dosage compensation complex on the X.

Co-amplified X/Y genes are highly expressed in testis and often have functions in chromosome segregation and RNAi, and we speculate that their parallel amplification on the X and Y is a result of ongoing X-Y interchromosomal conflicts over segregation. Sequence homology between putative drivers and their suppressors on the sex chromosomes, and their widespread targeting by small RNAs suggests that RNAi mechanisms are involved in silencing rampant sex chromosome drive. If amplified Y genes are involved in a battle with the X over fair transmission, changes in gene copy number may bias inclusion into functional sperm, and may trigger repeated co-amplification of distorters and suppressors on the sex chromosomes.

In principle, either sex chromosome could initiate this evolutionary tug-of-war over transmission into functional sperm. While we cannot determine the sequence of evolutionary events with certainty, the X chromosome is *a priori* more likely to acquire segregation distorters, creating strong selection to evolve suppressors on the Y. On one hand, natural selection is impaired on the non-recombining Y^3^, making drivers more likely to originate on the X. Additionally, the heterochromatic nature of a Y chromosome may render it especially vulnerable to be exploited by selfish elements during meiosis^46^.

Rampant sex chromosome drive can have important evolutionary consequences. Strong selective pressure to amplify Y-linked suppressors of meiotic drive may indirectly account for the complete genetic decay of the Y. Since the Y chromosome lacks recombination, strong positive selection for meiotic drive suppressors can propel linked deleterious mutations to fixation^17^, and the ongoing degeneration of ancestral Y genes may thus be a by-product of silencing recurrent meiotic drivers arising on the X. Patterns of molecular variation are suggestive of episodes of recurrent positive selection shaping neo-Y evolution of *D. miranda*^20^, and natural lines of *D. miranda* show a wide range of sex-ratio bias (with typically female-biased sex ratios^12^). These observations are consistent with recurrent and ongoing conflicts over segregation affecting the genomic architecture of sex chromosomes in this species.

Genetic conflict between X-Y ampliconic genes may also contribute to hybrid sterility and consequent reproductive isolation^33,47,48^. Segregation distortion can result in male hybrid sterility in Drosophila^49^, and further functional characterization of co-amplified, lineage-specific X-Y gene families will be needed to test the proposed link between X-Y genetic conflict and hybrid sterility.

X-Y interchromosomal conflict, and its consequent impact on gene amplification on sex chromosomes, may be widespread. In both human and mouse—two species with high-quality reference sequences for both sex chromosomes—the X and Y have co-acquired and amplified genes, and in both cases, meiotic drive has been invoked to explain this co-amplification^9,50–52,53^. Co-amplified genes have also been found in *D. melanogaster*^54^, and RNAi mechanisms have been shown to mediate suppression of sex ratio drive in flies^34–36^. Highly amplified gene families have been detected in other mammals^55^ and across fruit flies^56^, suggesting that sex chromosome drive may be prevalent in evolution; to determine the true phylogenetic range of lineage-specific acquisition and amplification of X-Y genes, high-quality sex chromosome assemblies across more taxa are needed.

## Materials and methods

### Genome and data availability

A 150-kb fragment of the Y chromosome was found to be missing in the previous genome assembly^14^ and the Y fragment was correctly reinserted before all downstream analyses. The updated genome assembly has been submitted to GenBank. All the data that were used and generated for this project are given in **Table S9**.

### *De novo* transcriptome assembly

To mask repeats in the genome, we used RepeatMasker^57^ with custom *de novo* repeat libraries, generated using RepeatModeler^58^ and Repdenovo^59^, along with the Drosophila repeat library from Repbase^60^. The *de novo* repeat libraries are given in **Data Supplement 1**, and a repeat-masked gff file is given in **Data Supplement 2**. Paired end RNA-seq reads from several male and female tissues (heads, carcass, whole body, testis, ovary, accessory gland, spermatheca, 3^rd^ instar larvae) were then aligned to the repeat-masked genome using HiSat2^61^ with the --dta parameter on default settings. The resulting alignment file was used to assemble the transcriptome using the software StringTie^62^ with default parameters. Fasta sequences of the transcripts were extracted from the gtf output produced by StringTie using the gffread utility.

### Gene annotation with Maker

We ran Maker^63^ three times to iteratively build and improve the gene annotation of the neo-sex chromosomes. For the first Maker run, we used annotated protein sequences for *D. melanogaster* and *D. pseudoobscura* from flybase.org, our *de novo* assembled *D. miranda* transcripts (see above), and the gene predictors Augustus^64^ and SNAP^65^ to get the initial set of predictions. The parameters est2genome and protein2genome were set to 1 to allow Maker to directly build gene models from the transcript and protein alignments, and we used the Augustus fly gene model and the SNAP *D. melanogaster* hmm file for this first run. The predictions from the first round were then used to train Augustus using BUSCO^66^ and also to train SNAP. The new Augustus gene model and SNAP hmm file were then used during the second Maker run, with the parameters est2genome and protein2genome set to 0. The maximum intron size was increased to 20000bp (default 10000bp). The results from the second round were then used to train Augustus and SNAP again, before the final round of Maker. This process resulted in a total of 21,524 annotated genes in *D. miranda*.

### Orthology detection

Transcript sequences for *D. pseudoobscura* were downloaded from flybase.org and only the largest transcript per gene was retained for downstream analyses. *De novo* annotated *D. miranda* transcripts were then aligned to this filtered *D. pseudoobscura* transcript set using BLAST^67^. Alignments with percentage identity <60% were discarded and the best alignment was calculated based on the e-value, score, % identity and alignment lengths. Each *D. miranda* transcript was thus assigned the ortholog that was its best BLAST hit. We identified paralogous genes in the *D. pseudoobscura* genome as those for which at least 80% of the sequence of one aligned to the other and vice versa. Paralogous genes in the *D. miranda* – *D. pseudoobscura* orthology calls were replaced by a single gene name from the duplicated gene family.

### Identifying multicopy genes

The gene annotation produced by Maker had roughly 2,500 more genes annotated on the Y/neo-Y compared to the neo-X, and hundreds of genes had multiple annotated copies on the Y/neo-Y chromosome (and also the X chromosome and autosomes in some cases). Based on the orthology calls from BLAST, 822 Maker annotated genes had more than 2 copies on the Y/neo-Y, and 209 of those genes had more than two copies on both the X/neo-X and Y/neo-Y. In our initial Maker annotation, 366 genes were missing on the neo-Y and 155 genes were missing on the neo-X. However, closer inspection revealed that the annotation was often fragmented, especially on the Y/neo-Y chromosome, which led to an overestimation of the number of distinct genes that had duplicated, but subsequent BLAST searches also revealed that Maker often failed to annotate individual copies of gene families. On the other hand, several genes in the annotation were “chimeras”, where two genes were collapsed into one by Maker and thus one of the genes appeared to be missing from the gene annotation, if it got assigned to the other *D. pseudoobscura* gene during orthology assignment. Thus, the actual number of missing genes is much smaller than our initial Maker annotation suggested. We thus manually verified, and if necessary fixed, each gene model that was annotated by Maker, and inferred to be either duplicated on the Y/neo-Y or missing from the neo-X and/or Y/neo-Y annotations. We used nucmer^18^ from the mummer package to individually align (one gene at a time) the sequences of their corresponding *D. pseudoobscura* orthologs to the *D. miranda* genome with the parameters --maxmatch and --nosimplify. Alignment coordinates were manually stitched together to get full gene coordinates. Only fragments that were at least 25% the length of the corresponding *D. pseudoobscura* ortholog, or at least 1000-bp long, were counted as duplicates/paralogs in the *D. miranda* genome. We also performed BLAST searches to identify the genes that had been lost from the neo-sex chromosomes.

In total, we annotate 6,448 genes on the neo-Y. Of these, 1,736 are ancestral single-copy Y genes (i.e. they were present on the ancestral autosome that formed the neo-Y). 1,105 of these genes were readily identified on both the neo-X and neo-Y by our Maker annotation, and are used as the single-copy orthology gene set in our analysis. 631 ancestral single-copy Y genes were initially missed or mis-qualified by our Maker annotation (i.e. 347 neo-Y genes were wrongly annotated as multi-copy by Maker, but our manual inspection revealed that they were present only as single-copy genes, and 114 neo-X genes and 170 neo-Y genes were missing from the Maker annotation, but found to be present on the neo-X and neo-Y, respectively, after manual checking using nucmer^18^.

In addition, we identify 457 genes (with 3,733 gene copies) that have become amplified on the neo-Y: 1,697 multi-copy Y genes (with a single copy on the neo-X), and 2,036 co-amplified neo-Y genes (which also amplified on the X/neo-X). In addition, we detect 959 genes in our Maker annotation that were not further considered. These “other” genes are comprised of 159 neo-Y genes that lack a homolog in *D. pseudoobscura*, 287 neo-Y genes that are present on an unknown location in *D. pseudoobscura*, 189 single-copy neo-Y genes that are present at multiple other locations in the genome (based on the Maker annotation), and 324 genes (from 49 unique proteins) with complicated mapping which could not be included in any categories of our analysis (i.e. genes for which the number of copies were ambiguous based on alignments such as for nested/overlapping genes; genes for which many alignments of variable identity were observed; genes which were amplifying on autosomes and the Y chromosome; chimeric genes).

Thus, after manual verification, we identified 94 genes that have co-amplified on the X/neo-X and the Y/neo-Y, with 647 copies on the X/neo-X and 2036 copies on the Y/neo-Y (and 58 copies on the autosomes). We also identified 363 genes that have only amplified on the Y/neo-Y chromosome, with a total of 1697 copies on the Y/neo-Y. Thus, the Y/neo-Y chromosome has gained at least 3200 gene copies.

We identified 17 genes that are present on chr3 in *D. pseudoobscura* but are missing on the neo-X in *D. miranda*; 6 of those genes are found on other chromosomes in the *D. miranda* genome and 6 are still present on the Y chromosome in *D. miranda*. We identified 143 genes that are present on chr3 in *D. pseudoobscura* but absent on the neo-Y in *D. miranda* and 138 of those are still present on the neo-X in *D. miranda*. However 5 genes have been lost from both the neo-X and neo-Y chromosomes and BLAST searches failed to identify other chromosomal locations that those genes could have moved to. Genome annotations for all genes, multi-copy Y genes and their homologs, and co-amplified X and Y genes are given in **Data Supplements 3-5**.

Karyotype plots showing co-amplified X/Y genes were produced using karyoploteR^68^. Plots showing multi-copy gene locations on the neo-X and neo-Y (Y1 and Y2) were created using genoPlotR^69^.

### Y-chromosome replacement lines

Y chromosomes from seven *D. miranda* lines (**Table S7**) were moved into an MSH22 (reference) background by repeated backcrossing of hybrid males (8 generations) with virgin MSH22 females. We then extracted DNA from a single male from each Y-chromosome replacement line using a Qiagen DNeasy kit following the standard extraction protocol. DNA libraries were prepared using the Illumina TruSeq Nano Prep kit and sequenced on a Hiseq 4000 with 100bp PE reads. Raw reads from these seven Y-chromosome replacement lines along with sequencing data for three isofemales lines (MA03.4, 0101.7, MA03.2) were initially mapped to the reference MSH22 genome using BWA mem^70^. The resulting files were processed using Samtools^71^ and PCR duplicates were removed using Picard Tools. We called SNPs using GATK’s UnifiedGenotyper^72^ and filtered SNPs using VCFtools^73^ and retained biallelic SNPs, positions with no more than 50% of individuals missing a call, and individual genotypes with GQ > 30 and depth > 3 and < 80.

To confirm Y replacement, we first estimated nucleotide diversity (π) using VCFtools across each chromosome with the expectation that it should be uniformly low given that MSH22 is an inbred line. We noted a few regions that showed peaks of elevated heterozygosity (**Figure S5A**), which is indicative of either residual heterozygous regions in the MSH22 line or suggests that our backcrossing failed to replace all chromosomes with MSH22 chromosomes. Further, the MSH22 Y^BB51^ line appeared to be heterozygous across all of chr4 while all its other chromosomes appeared to be replaced with MSH22 chromosomes. However, given that variation on chr4 of line BB51 likely contributes little to Y chromosome gene amplification estimates, we retained MSH22 Y^BB51^ in our analyses. Phylogenetic networks using SplitsTree^74^ for chr2 (17,189 SNPs) and the Y chromosome (1,543 SNPs) confirmed that chr2 of Y-chromosome replacement lines appeared identical to MSH22 while the seven Y chromosomes all appeared genetically distinct (**Figure S5B, C**).

### Read coverage analysis to infer gene copy number

We used DIAMOND^75^ to align raw Illumina reads from each Y-chromosome replacement line to the longest isoform for each *D. miranda* protein (n=12,180). Only the top hit for each read was retained, and mean coverage over each protein was estimated using Bedtools^76^. To estimate the copy number for co-amplified Y, multi-copy Y, and multi-copy autosome and X genes we normalized estimates using median coverage over 98 randomly selected X-linked single copy genes.

### Phylogenetic analysis of co-amplified X/Y genes

Co-amplified X/Y gene regions and their *D. pseudoobscura* ortholog were aligned using MAFFT^77^. Due to the fragmented nature of some Y copies, a small number of copies were removed to maximize the number of informative sites while retaining most gene copies. We created rooted maximum-likelihood phylogenetic trees using RAxML 8.2.9^78^ with 200 bootstrap replicates and a GTR + gamma model of sequence evolution. Phylogenetic trees were visualized using FigTree version 1.4.3 (https://github.com/rambaut/figtree/).

### Gene expression analysis

Kallisto^79^ was used to quantify gene expression and calculate TPM values for each gene in our annotation using several male and female tissues (whole body, carcass, 3^rd^ instar larvae, gonads, spermatheca and accessory gland) using default parameters and 100 bootstraps. The R function heatmap.2 from the gplots package (https://cran.r-project.org/web/packages/gplots/index.html) was used to plot heatmaps to visualize tissue-specific differences in gene expression. Each row in the heatmap is a different gene and the different columns represent different tissues. The heatmap was row-normalized (each row was scaled to have mean 0 and standard deviation 1) to indicate the tissue with the highest expression for each gene. The tissue-specificity index, *τ* was calculated in the following manner:

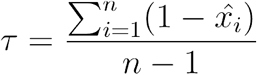

where, *x*_*i*_ is the expression of the gene (TPM) in tissue i, n is the number of tissues, and

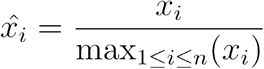

### GO analysis

GO analysis was done using GOrilla^80^ and the *D. melanogaster* orthologs of the genes co-amplifying on the X and Y was used as the target set. The GO terms that were enriched and had a p-value less than 10^−3^ and fdr less than 0.05 were visualized using the software Revigo^81^.

### Testis RNA libraries

We dissected testes from 3-8 day old virgin males of *D. miranda* (strain MSH22) reared at 18°C on Bloomington food. We used Trizol (Invitrogen) and GlycoBlue (Invitrogen) to extract and isolate total RNA. We resolved 20 µg of total RNA on a 15% TBE-Urea gel (Invitrogen) and size selected 19-29 nt long RNA, and used Illumina’s TruSeq Small RNA Library Preparation Kit to prepare small RNA libraries, which were sequenced on an Illumina HiSeq 4000 at 50 nt read length (single-end). We used to Ribo-Zero to deplete ribosomal RNA from total RNA, and used Illumina’s TruSeq Stranded Total RNA Library Preparation Kit to prepare stranded testis RNA libraries, which were sequenced on an Illumina HiSeq 4000 at 100 nt read length (paired-end).

### Analysis of testes smRNA and testes totalRNA data

Stranded total RNA paired-end reads were mapped to the *D. miranda* genome using HiSat2^61^ with default parameters and the --rna-strandness parameter set to RF. Single-end small RNA-seq reads were aligned to the genome using bowtie2^82^ and default parameters. BamCoverage from the deeptools package^83^ was used to convert bam alignment files to bigwig format in both cases to be visualized using IGV. Sense and antisense transcription estimates were obtained based on the alignment and the orientation of genes using bedtools^84^. The number of small RNA and total RNA reads mapping to the co-amplified X/Y genes were summed for each gene family, based on sense or antisense transcription, and barplots of counts were plotted in log2 scale using R (Figure 5D and 5E). The number of small RNA reads mapping to each annotated gene (sense and antisense counts) were divided by the gene/fragment length and boxplots were plotted in R for single copy Y genes, genes that have only amplified on the Y and genes that have co-amplified on the X and Y (Figure 5G).

## Acknowledgements

Funded by NIH grants (R01GM076007, GM101255 and R01GM093182) to DB. We thank L. Gibilisco for generating small RNA libraries and K. Chatla and A. Tran for generating genomic libraries.

## Data availability

BioProject ID PRJNA545539 and Dryad.

---

**Table S1.** Gene loss on the different Muller elements of *D. miranda.* Shown are genes in *D. pseudoobscura* for different chromosomes that are absent on their homologous chromosome arm in *D. miranda*. The table gives *D. pseudoobscura* gene ID (FBgn), *D. melanogaster* ortholog, the chromosomal location of the gene in *D. pseudoobscura*, the tissue of highest expression in *D. pseudoobscura*, the category of the gene, and whether the gene is present somewhere else in the *D. miranda* genome.

**Table S2.** Overview of multi-copy Y genes. Shown are total copy numbers for multi-copy Y genes, as well as the number of full-length (>90%) and partial Y copies (50-90%; 25-50%; and less than 25% compared to the length of the orthologous gene in *D. pseudoobscura*). The expression spreadsheet shows expression of orthologs of multi-copy Y genes in *D. pseudoobscura*, and the GO analysis shows the orthologous genes in *D. melanogaster*. No significant GO enrichment terms were detected.

**Table S3.** Overview of co-amplified X/Y genes.. Shown are total copy number for co-amplified X and Y genes, as well as the numbers of full-length (>90%) and partial X and Y copies (50-90%; 25-50%; and less than 25% compared to the length of the orthologous gene in *D. pseudoobscura*). The expression spreadsheets show expression of orthologs of co-amplified X/Y genes in *D. pseudoobscura* and *D. melanogaster*, and GO analysis shows the orthologous genes in *D. melanogaster* and GO terms that were significantly enriched among co-amplified X/Y genes (using either Gorilla or PantherDB).

**Table S4.** Expression of individual copies of multi-copy Y genes and co-amplified X/Y genes. Shown is the inferred fraction of individual gene copies expressed, depending on different cut-offs.

**Table S5.** Genome intervals of clustered (within 100kb of each other) multi-copy genes and their *Drosophila pseudoobscura* gene ID (FBgn).

**Table S6.** Genome intervals of clustered (within 100kb of each other) co-amplified genes and their *Drosophila pseudoobscura* gene ID (FBgn).

**Table S7.** Y-chromosome replacement lines used in the study. Shown is the collection location for the different Y chromosome replacement lines generated.

**Table S8.** Small RNA mapping from testis, for single copy X and Y genes, multi-copy Y genes and their X homologs, and co-amplified X and Y genes

**Table S9.** Sequence data generated and SRA accession numbers.

**Figure S1. Validation of multi-copy and co-amplified Y genes in the *D. miranda* genome assembly using Illumina read coverage analysis from the reference strain (MSH22).** Shown is the gene copy number from the genome assembly and the estimated copy number based on read mapping to A) multi-copy genes (Pearson correlation = 0.82) and B) co-amplified genes (Pearson correlation = 0.84) with the 1:1 line shown. Zoomed in regions for multi-copy and co-amplified genes are shown in C) and D). Note that our coverage analysis typically underestimates the number of Y-linked copies found in the assembly, presumably due to many multi-copy genes being fragmented in the assembly.

**Figure S2. Location of multi-copy Y genes.** The genomic coordinates of genes identified as multi-copy on the two largest Y chromosome scaffolds (Y1 and Y2) and their corresponding single copy location on the neo-X (Muller C). Most multi-copy genes on the Y exist in tandem and therefore there is a substantial overlap of plotting lines. Note that the Y/neo-Y chromosomes in our current assembly in *D. miranda* consists of three major scaffolds (Y1, Y2, Y3, with Y3 possibly corresponding to the ancestral Y).

**Figure S3. Location of orthologs of multi-copy Y genes and co-amplified X/Y genes in *D. pseudoobscura* and *D. miranda*.** Each line shows the location of the ortholog of a co-amplified X/Y gene (in blue), or a multi-copy Y gene (in orange) along *D. pseudoobscura* chr3, or the neo-X of *D. miranda*. Of the 363 multi-copy genes identified on the Y/neo-Y of *D. miranda*, 349 are located on chr3 of *D. pseudoobscura*, 2 genes are on chr2, 7 genes with unknown location, 1 gene on XL, and 4 genes on XR. Of the 94 co-amplified X/Y genes, 59 are located on chr3 of *D. pseudoobscura*, 4 genes are on chr2, 1 gene on chr4, 16 genes with unknown location, 6 genes on XL, and 8 genes on XR.

**Figure S4. Phylogenetic relationships of co-amplified genes in *D. miranda***. Maximum-likelihood trees of *D. miranda* X and Y gene copies with nodes showing >70 bootstrap support highlighted with black circles. X-linked copies are shown in red, Y-linked copies shown in blue, with distinct X and Y groupings collapsed. Fasta alignments are in **Data Supplement 15**.

**Figure S5. Properties of Y-chromosome replacement lines.** A) Individual nucleotide diversity (pi) shown for each replacement line. Peaks identify regions of residual heterozygosity in each Y-chromosome replacement line. B) Phylogenetic networks of autosomal SNPs of the seven Y-chromosome replacement lines and three lines from which some Y chromosomes were derived. As expected, all Y replacement lines cluster with MSH22. C) Phylogenetic network of SNPs showing seven distinct Y types.

**Figure S6.** Copy number estimates for co-amplified Y genes, multi-copy Y genes, and multi-copy autosome and X genes. For co-amplified Y genes we show all genes that were identified as co-amplified. For the multi-copy Y genes we only show genes with >3 copies on the Y. For multi-copy autosome and X genes we show only genes with >4 total copies. Multi-copy autosome and X estimates are predicted to be highly similar given that the autosome and X background in each Y-chromosome replacement line is nearly identical. Slight deviations are likely due stochasticity in sequencing and read mapping, residual heterozygosity in the MSH22 line, or unique Y chromosome gene amplifications.

**Figure S7**. Repeats may contribute to accumulation of multi-copy genes. Transposable elements often flank multi-copy Y genes, and may have contributed to amplification of genes on the X and Y. Genes are shown in green, and TEs are shown in gray.

**Figure S8.** Expression patterns of X- and Y-linked genes for **A.** single-copy X- and Y-linked genes; **B.** multi-copy Y-linked genes and their X homologs; **C.** co-amplified X and Y genes. Expression for individual gene copies is shown. Expression values were row-normalized to obtain Z-scores with a mean of 0 and standard deviation of 1, using the built-in scale = ‘row’ argument in the heatmap.2 function from the package gplots in R. The data shown are presented in **Data Supplement 16**.

**Figure S9**. Expression patterns of X- and Y-linked genes for **A.** single-copy X- and Y-linked genes; **B.** multi-copy Y-linked genes and their X homologs; **C.** co-amplified X and Y genes. Total summed expression for all gene copies of a gene family is shown. Expression values were row-normalized to obtain Z-scores with a mean of 0 and standard deviation of 1, using the built-in scale = ‘row’ argument in the heatmap.2 function from the package gplots in R. The data shown are presented in **Data Supplement 17**.

**Figure S10.** Tissues-specific expression patterns of orthologs of co-amplified X/Y genes in *D. melanogaster*. FPKM values were downloaded from flybase.org. The data shown are presented in **Data Supplement 18**.

## Data Supplements (submitted on dryad)

Data Supplement 1. Repeat library used for masking the *D. miranda* genome (fasta file).

Data Supplement 2 Repeat annotation of the *D. miranda* genome (gff file).

Data Supplement 3 Gene annotation (all genes) of the *D. miranda* genome (gff file).

Data Supplement 4 Gene annotation of multi-copy Y genes and their orthologs in the *D. miranda* genome (gff file).

Data Supplement 5 Gene annotation of co-amplified X and Y genes in the *D. miranda* genome (gff file).

Data Supplement 6. Gene expression values (TPM) for multi-copy Y genes, and co-amplified Y genes, in different *Drosophila miranda* male tissues, and tissue-specificity index tau.

Data Supplement 7. MSL ChIP and Input counts (normalized to library size) for neo-X genes whose homolog is classified as single-copy or multi-copy Y/neo-Y.

Data Supplement 8. Gene copy numbers for co-amplified X/Y genes.

Data Supplement 9. Gene expression values (TPM) for co-amplified X and Y gene families, in different *Drosophila miranda* male tissues.

Data Supplement 10. Gene expression values (FPKM) for orthologs of co-amplified X/Y genes in different *Drosophila pseudoobscura* male and female tissues.

Data Supplement 11. Sense and anti-sense testis total RNA summed counts for co-amplified genes.

Data Supplement 12. Sense and anti-sense testis small RNA summed counts for co-amplified genes.

Data Supplement 13. Testis total RNA counts for all copies of a gene family for different categories of genes on the X/neo-X and Y/neo-Y chromosome.

Data Supplement 14. Testis small RNA raw counts for different categories of genes on the X/neo-X and Y/neo-Y chromosome.

Data Supplement 15. Fasta alignment of co-amplified X/Y genes and their *D. pseudoobscura* ortholog.

Data Supplement 16. Gene expression values (TPM) for single-copy Y genes, multi-copy Y genes, and co-amplified Y genes, and their neo-X/X homologs in different *Drosophila miranda* tissues. Expression values for all copies of a gene family are shown individually.

Data Supplement 17. Gene expression values (TPM) for multi-copy Y genes, and co-amplified X and Y genes in different *Drosophila miranda* tissues. Expression values for all copies of a gene family are summed.

Data Supplement 18. Gene expression values (FPKM) for orthologs of co-amplified X/Y genes in different *Drosophila melanogaster* tissues.

